# Whole-brain modelling of resting state fMRI differentiates ADHD subtypes and facilitates stratified neuro-stimulation therapy

**DOI:** 10.1101/2020.05.08.084491

**Authors:** Behzad Iravani, Artin Arshamian, Peter Fransson, Neda Kaboodvand

## Abstract

Recent advances in non-linear computational and dynamical modelling have opened up the possibility to parametrize dynamic neural mechanisms that drive complex behavior. Importantly, building models of neuronal processes is of key importance to fully understand disorders of the brain as it may provide a quantitative platform that is capable of binding multiple neurophysiological processes to phenotype profiles. In this study, we apply a newly developed adaptive frequency-based model of whole-brain oscillations to resting-state fMRI data acquired from healthy controls and a cohort of attention deficit hyperactivity disorder (ADHD) subjects. As expected, we found that healthy control subjects differed from ADHD in terms of attractor dynamics. However, we also found a marked dichotomy in neural dynamics within the ADHD cohort. Next, we classified the ADHD group according to the level of distance of each individual’s empirical network from the two model-based simulated networks. Critically, the model was mirrored in the empirical behavior data with the two ADHD subgroups displaying distinct behavioral phenotypes related to emotional instability (i.e., depression and hypomanic personality traits). Finally, we investigated the applicability and feasibility of our whole-brain model in a therapeutic setting by conducting in silico excitatory stimulations to parsimoniously mimic clinical neuro-stimulation paradigms in ADHD. We tested the effect of stimulating any individual brain region on the key network measures derived from the simulated brain network and its contribution in rectifying the brain dynamics to that of the healthy brain, separately for each ADHD subgroup. This showed that this was indeed possible for both subgroups. However, the current effect sizes were small suggesting that the stimulation protocol needs to be tailored at the individual level. These findings demonstrate the potential of this new modelling framework to unveil hidden neurophysiological profiles and establish tailored clinical interventions.

## INTRODUCTION

The recent advancements in developing and applying large-scale, non-linear brain modelling methods to neuroimaging data have proven to be highly efficient in explaining, predicting and integrating neuronal activity with complex behavior (Breakspear 2017; McIntosh and Jirsa 2019). Importantly, these methodological developments have enabled us to discover parameters related to brain dynamics that are otherwise not directly observable but have been shown to drive complex behavior (Breakspear 2017; McIntosh and Jirsa 2019; Kaboodvand 2019). This has been especially evident in the quest to understand the underlying mechanisms for a wide range of brain disorders (e.g., epilepsy; (Jirsa et al. 2014)) with a potential to offer insights on how to design therapeutic interventions. Here, we do both by first applying a novel type of adaptive frequency-based modelling of whole-brain oscillations (Kaboodvand et al. 2019) on Attention Deficit Hyperactivity Disorder (ADHD) to disclose empirically unknown, but theoretically supported, functional network subtypes of ADHD, and second by providing the first insights in how to modulate these oscillations, enabling optimization of neuro-stimulation protocols for different subgroups within each disorder, so-called stratified neuro-stimulation therapy.

ADHD is one of the most common neurodevelopmental disorders affecting children with symptoms starting in early childhood but that in many cases continue into adulthood (Petrovic and Castellanos 2016) affecting approximately 3.4% of the worldwide adult population (Fayyad et al. 2007). ADHD is associated with a wide variety of symptoms such as lack of sustaining attention, and/or impulsivity and hyperactivity (Biederman et al. 2010; Castellanos and Proal 2012; De La Fuente et al. 2013; Kaboodvand 2019). Critically, ADHD symptoms are hard to disentangle from other maladies, such as depression (Wilens et al. 2002) and bipolar disorder (Youngstrom et al. 2010). Importantly, although ADHD traditionally has been considered as a dysfunctional regulation of non-emotional information processing, recent insights from both theoretical (Petrovic and Castellanos 2016), as well empirical data from brain anatomy (Bayard et al. 2020) support the notion that ADHD and disorders involving emotional instability such as conduct disorder share many traits. Thus, many of the observed emotional subtypes of ADHD may not always result from comorbidity, but should be classified using a dimensional approach to psychiatric disorders (Petrovic and Castellanos 2016). This has, however, been hard to achieve as both real and apparent co-morbidity together with the heterogeneous nature of ADHD have hampered research to find robust biomarkers, and thus enable tailored interventions across the ADHD spectrum (Rubio et al. 2016; Wåhlstedt et al. 2009).

Previous studies have found ADHD-related changes in brain function (Krain and Castellanos 2006; Wilens and Spencer 2010; Kaboodvand et al. 2020; Kaboodvand 2019) and gray matter structure (Batty et al. 2010; Carmona et al. 2005), nevertheless, no significant changes in the white matter structure have been reported (Batty et al. 2010; Carmona et al. 2005). Moreover, changes both in brain structure and function in ADHD have previously been suggested to be a disorder of attractor dynamics in computational studies of ADHD (Durstewitz et al. 2020; Hauser et al. 2016). However, to the best of our knowledge, there is no previous study that supports this hypothesis. Importantly, a dynamical system modeling framework permits the investigator to scale down biophysical and structural processes of relevance to the study disease in the brain to a low-dimensional manifold, including attractors to model dynamic phenomenon as attractor states as well as allowing for modelling of the influence from behavioral (e.g. distractibility and emotional stability) variables, which can be parametrized and compared across cohorts (Kaboodvand 2019; Durstewitz et al. 2020).

In the present study, first, we apply a novel adaptive frequency-based modelling of whole-brain oscillations (Kaboodvand et al. 2019) to resting-state fMRI data collected in a control and ADHD cohort with the aim to disclose ADHD subtypes in the context of large-scale network modelling that has been previously glossed over in empirical investigations but that are supported by theoretical arguments. In essence, our whole-brain model consists of nonlinear differential equations that are coupled together in accordance to the connectivity patterns provided by structural MRI connectivity data. Indeed, anatomical and functional neuroimaging techniques have proven useful yet under-used in clinical diagnosis and therapy. Whole-brain dynamic modeling helps us integrate multimodal neuroimaging data and bridge the gaps between different levels of observation and analysis (Stefanovski et al. 2016). Further, it provides a platform to understand the fundamental mechanisms of brain function in health and disease, as well as to study the intervention effects (Durstewitz et al. 2020; Kaboodvand 2019). Characterization of whole-brain dynamics as macroscopic phenomena provides a holistic view which is robust with respect to interference from modest levels of system noise, e.g. head-motion and inter-individual signal variance. We hypothesized that the ADHD cohort should exhibit more unstable patterns of dynamics as well as a different involvement of brain structures compared to controls that will enable a potential categorization of ADHD subtypes.

Second, based on the results obtained in the first step, we aimed to provide insights on how wholebrain dynamic modelling can be used to find robust objective targets for designing therapeutic interventions and controlling individual responses to brain stimulation via designing goal-directed neuro-stimulation protocols (Deco and Kringelbach 2014; Kaboodvand et al. 2020; Durstewitz et al. 2020; Kaboodvand et al. 2019). To this end, we used an *in silico* excitation simulation protocol to shift the dynamics of ADHD resting-state networks to that of healthy controls. This simulation was performed by mimicking a clinical neuro-stimulation (e.g. transcranial magnetic stimulation (TMS)) effect on different brain regions and assessed the possibility to shift the dynamic regime of the targeted brain region from a stochastic fluctuations’ domain into an oscillatory domain by changing the local bifurcation parameter of that region and subsequently comparing the network measures derived from simulated static functional connectivity (FC) network of ADHD and healthy controls to evaluate the therapeutic effect of every stimulation target (Kaboodvand et al. 2019). Applying repetitive TMS has been shown to selectively change the oscillatory activity of the targeted region and induce various local neural effects (Okamura et al. 2001; Thut and Pascual-Leone 2010); yet we only simulated the effect of inducing oscillatory activity in different targeted regions, to further elaborate the heterogeneity within an ADHD cohort and highlight the importance of designing stratified neuro-stimulation protocols.

## METHOD

### Dataset and preprocessing

The primary data source for this study included two open access datasets. First, we used the Human Connectome (HCP) 500 subject release (Van Essen et al. 2013; Smith et al. 2013) to model wholebrain dynamics in healthy individuals (Control). We did this by using a densely sampled structural and functional scaffold, that included resting-state fMRI (rs-fMRI) and diffusion tensor imaging (DTI) data as well as the T1-weighted images of the same individuals (n = 447, age range: 21-50 yr). The rs-fMRI data were collected during 14.4 minutes with the temporal resolution of 0.72 seconds (1200 image volumes). Further information regarding the MR acquisition parameters can be found in (Van Essen et al. 2013). Briefly, preprocessing of rs-fMRI data included removing nuisance covariates (global signal, mean white matter and cerebrospinal fluid signals), linear and quadratic trends, followed by band-pass filtering (0.02–0.12 Hz). Further details on data preprocessing can be found in (Kaboodvand et al. 2019). For the ADHD cohort, we used resting-state fMRI data from subjects diagnosed with ADHD (n = 40, age range: 21-50 yr) provided by the University of California LA Consortium for Neuropsychiatric Phenomics study (Poldrack et al. 2016). Identical data preprocessing steps were carried out for the ADHD resting-state data as for the healthy cohort.

We used the Desikan-Killiany parcellation of the cortex into 68 regions of interest (ROIs) (Desikan et al. 2006), combined with 14 subcortical regions derived from FreeSurfer segmentation (Makris et al. 2008; Seidman et al. 1997). High-quality diffusion-weighted MRI data for the same 500 subjects from the HCP consortium (Van Essen et al. 2012; Glasser et al. 2013) was used for a streamline tractography. Next, we created subject-level weighted structural connectomes using measures of streamline density, computed by dividing the number of streamlines connecting two regions by the average of the volumes of the two interconnected regions (van den Heuvel et al. 2012). For details on the processing steps of diffusion-MRI derived connectivity data, we refer the reader to (de Reus and van den Heuvel 2014). We then constructed a group-representative structural connectome by averaging the subject-level structural connectivity (SC) entries which had nonzero values for at least 60% of the subjects (de Reus and van den Heuvel 2013), followed by resampling the data to follow a Gaussian distribution with μ= 0.5 and σ= 0.15 (van den Heuvel et al. 2015).

### Computational modelling of brain dynamics

Our model is constructed from weakly coupled oscillators which are interacting with each other through the structural connectome. Each oscillator represents a brain region with two coupled differential equations as follows:

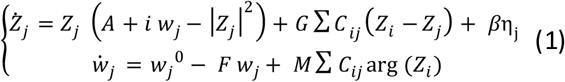

The model parameters of interest here are G (scalar, global coupling), M (scalar, modulation index), C (a matrix that contains the degree of SC between all brain regions), F (scalar, feedback coefficient) and A (scalar, global bifurcation parameter). The state variables are Z and w where Z is complex-valued BOLD-like signal with time-varying intrinsic frequency (w). The control parameters G and M regulate the amount of the interaction between different oscillators with larger values of G and M implying a stronger coupling. Model parameters A and F govern the local dynamics of oscillators, where a larger value of A is associated with higher amplitude and larger F is tied to faster convergence to steady state intrinsic frequency. For further details please see (Kaboodvand et al. 2019). Furthermore, additive Gaussian noise with a standard deviation of β= 0.02, implemented as a Wiener process, was added to the model. The output of the model is a set of complex-valued BOLD-like signals (Z) with time-varying intrinsic frequencies (w). The real part of Z is considered to represent BOLD signal recorded in the fMRI experiments whereas the imaginary part characterizes the hidden state of the oscillator that it is unseen to the scanner.

### Parameter optimization

In order to find the optimal working point of the model, we need to find the parameter set which maximizes the similarity between empirical and simulated BOLD signals (covering 82 ROIs). Accordingly, the set of model-parameters, including global coupling, bifurcation, modulation index and the feedback coefficient (G, A, M and F) were explored in a grid-search framework to find the optimal model working point in the 4-dimensioned parameter space. This was done separately for the Control and ADHD data cohorts. Notably, for the whole-brain modelling approach we used the SC derived from the high quality and large sample size HCP 500-subject release, given that previous studies have shown that the SC is not significantly different between healthy and ADHD cohorts (Batty et al. 2010; Carmona et al. 2005). Initially, the healthy control model was fitted to the HCP dataset and subsequently further validated against the control UCLA dataset. Further information on this validation step and the results thereof can be found in Supplementary Figure S1. Importantly, given the parameter set derived from fitting the Healthy model to the HCP dataset, the correlation between the simulated FC and the empirical FC obtained from the UCLA dataset was even larger (rho = 0.40) than the correlation between the simulated FC and the empirical FC obtained from the HCP dataset (rho = 0.34) (Supplementary Figure S1).

Our search space spanned A from −0.07 to 0.07 with a step-size of 0.002, G ranged from 0.002 to 0.03 with a step-size of 0.002, M ranged from −0.6 to −0.2 with a step-size of 0.1, and F covered the range of 0.1 to 0.2, with a step-size of 0.01 resulting in total of 58575 simulations.

It is worth mentioning that for each point in the parameter space, the real part of system variables (i.e., Z) was used as the simulated BOLD signals for all 82 ROIs. The model simulated the acquisition of 6 minutes resting-state BOLD data, similar to the empirical data. To assess the model performance, the FC of resting state networks were compared between simulated and empirical data. Estimation of FC was performed through standard FC analysis of resting state fMRI; started by band-pass filtering of empirical BOLD (0.02-0.12 Hz), following by Pearson correlation and fisher z-transformation to estimate simulated and empirical FC. Next, the Pearson correlation coefficient was obtained between the FC matrices derived from empirical and simulated data using 500 permutations Monte Carlo sampling approach.

In Monte Carlo simulation, for each of the 500 iterations, same procedure as mentioned above was performed in which the matrices of correlation coefficients were computed from the pre-processed empirical and simulated BOLD signals originating from 82 ROIs as described above. Subsequently, we constructed the group-level representation for that specific iteration by averaging the subject-level FC matrices (only half of the dataset was randomly selected and used in each iteration). The same procedure was applied to the simulated BOLD signals to create a simulated FC matrix for each set of parameters (**Figure 1**).

**Figure 1.**
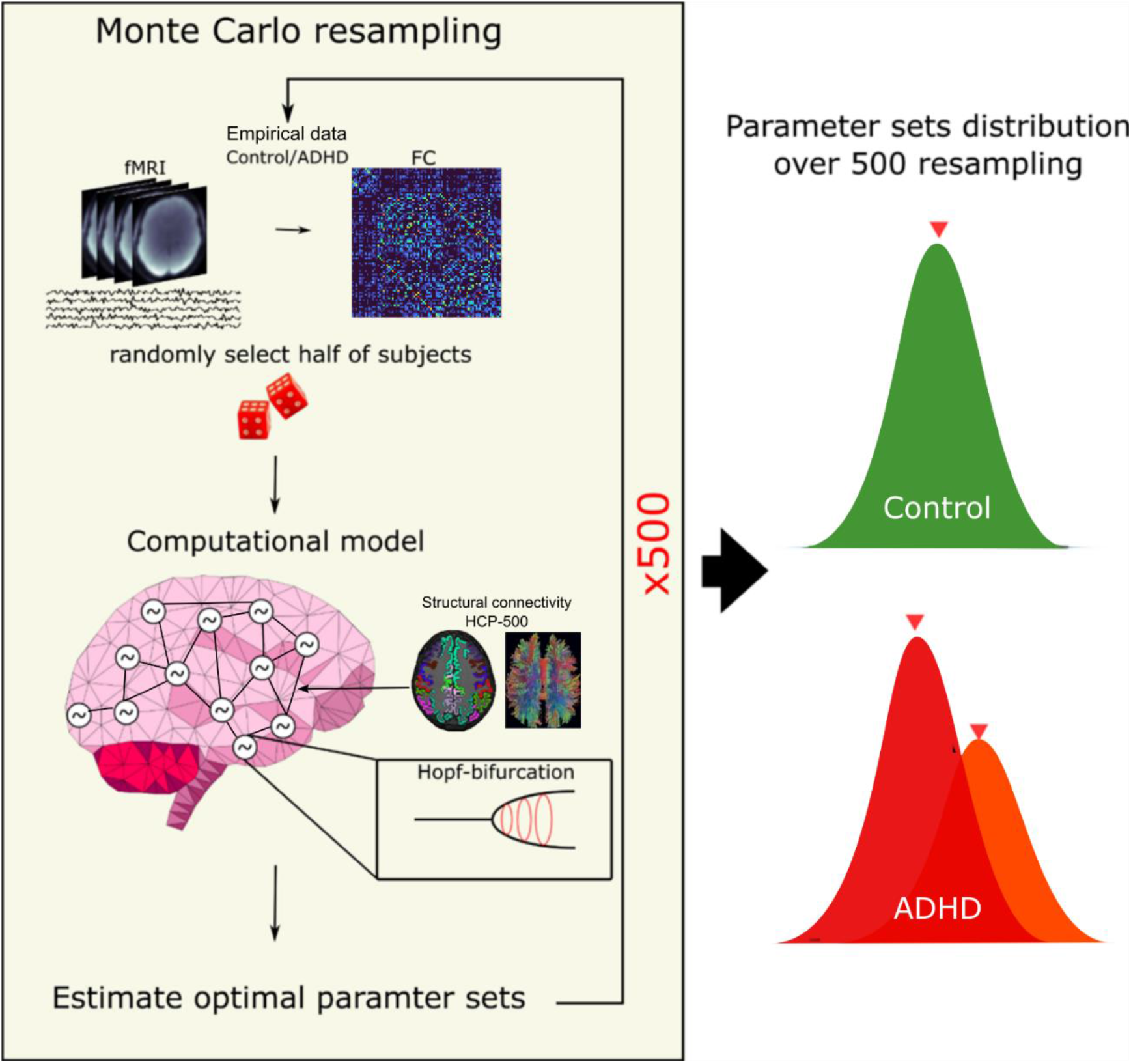
Parameter optimization using Monte Carlo simulations. Half of the data were randomly selected, and the corresponding functional connectivity was used to optimize the model parameters, repeated 500 times. The aggregated distribution (i.e. G, A, M and F) of estimated parameters across 500 iterations was examined, revealing bimodal distribution for ADHD.

Finally, we computed the Pearson correlation coefficient between the empirical FC matrix and the FC matrix achieved for the simulated signals as a measure of similarity between the empirical and simulated BOLD signals. After these 500 iterations, a histogram of parameters was obtained, and we estimated the most likely parameter set for each cohort that maximized the similarity between simulated and empirical data.

### Modelling focal brain stimulation by a nodal excitation protocol *in silico*

To address the question of to which extent it is possible to change the regime of brain dynamics by local stimulation, we applied a protocol of excitatory input to our model that served to mimic focal brain stimulation. The local dynamics of oscillators in our whole-brain model are working close to the edge of Hopf bifurcation (i.e. bifurcation parameter A was close to zero). We induced a local excitatory effect of focal brain stimulation by shifting the dynamical regime of a region via moving its bifurcation parameter from the optimal parameter value to 1, where it surpasses the Hopf birfucation and starts oscillating, resembling a focal brain stimulation (e.g. TMS effect). Bifurcation-induced excitatory stimulation was applied to all possible targets (82 ROIs). To assess the effect of local excitatory stimuli for all brain regions in our model, we computed for each node three network centrality measures: clustering coefficient, nodal efficiency and nodal strength for the simulated resting state network. Next, network measures for all 82 nodes were combined to a common feature space with 246 dimensions (82×3). The effect of stimulating each region was measured by computing the pairwise Euclidean distance of the feature vectors (i.e. network centrality measures) between the ADHD and Control model.

## RESULTS

### Optimal model parameters for the ADHD and Control cohort

We used a whole-brain, adaptive frequency-based weakly coupled oscillators model (Kaboodvand et al. 2019) to assess putative differences of brain dynamics in ADHD compared to a Control group. In attempt to find robust parameter sets we used the Monte Carlo simulation technique for parameter optimization of our dynamical model. This enabled us to estimate the most likely parameter sets that best explained the empirical data for both Control and ADHD cohorts (**Figure 1**).

The distribution of the parameter sets derived from 500 iterations for the ADHD cohort showed two local maximums, but it was a unimodal distribution of the Control cohort (see also schematic in **Figure 1**). Accordingly, we divided the ADHD cohort into two subgroups based on the level of Euclidian distance of every individual empirical FC network from the model-based simulated network, here called ADHD1 and ADHD2, with 25 and 15 subjects in each group, respectively. Particularly, we computed the Euclidian distance between each patient’s empirical FC matrix from the two model-derived simulated FC matrices, to find which model better explains the empirical data for each patient. Accordingly, we could designate each patient to one of the two ADHD subgroups, associated with the two ADHD models (i.e. ADHD1 or ADHD2). Next, we assessed the similarity between simulated FC from the computational model and empirical data for all three cohorts namely Control, ADHD1 and ADHD2 (rho = 0.34, 0.43 and 0.41 respectively; **Figure 2A**). Of note, compared with the results shown in our previous study (Kaboodvand et al., 2019) the degree of similarity between wholebrain based connectivity for the simulated datasets and the whole-brain connectivity observed for the empirical signals is lower which may be due to inclusion of 14 additional subcortical regions in the present study.

**Figure 2.**
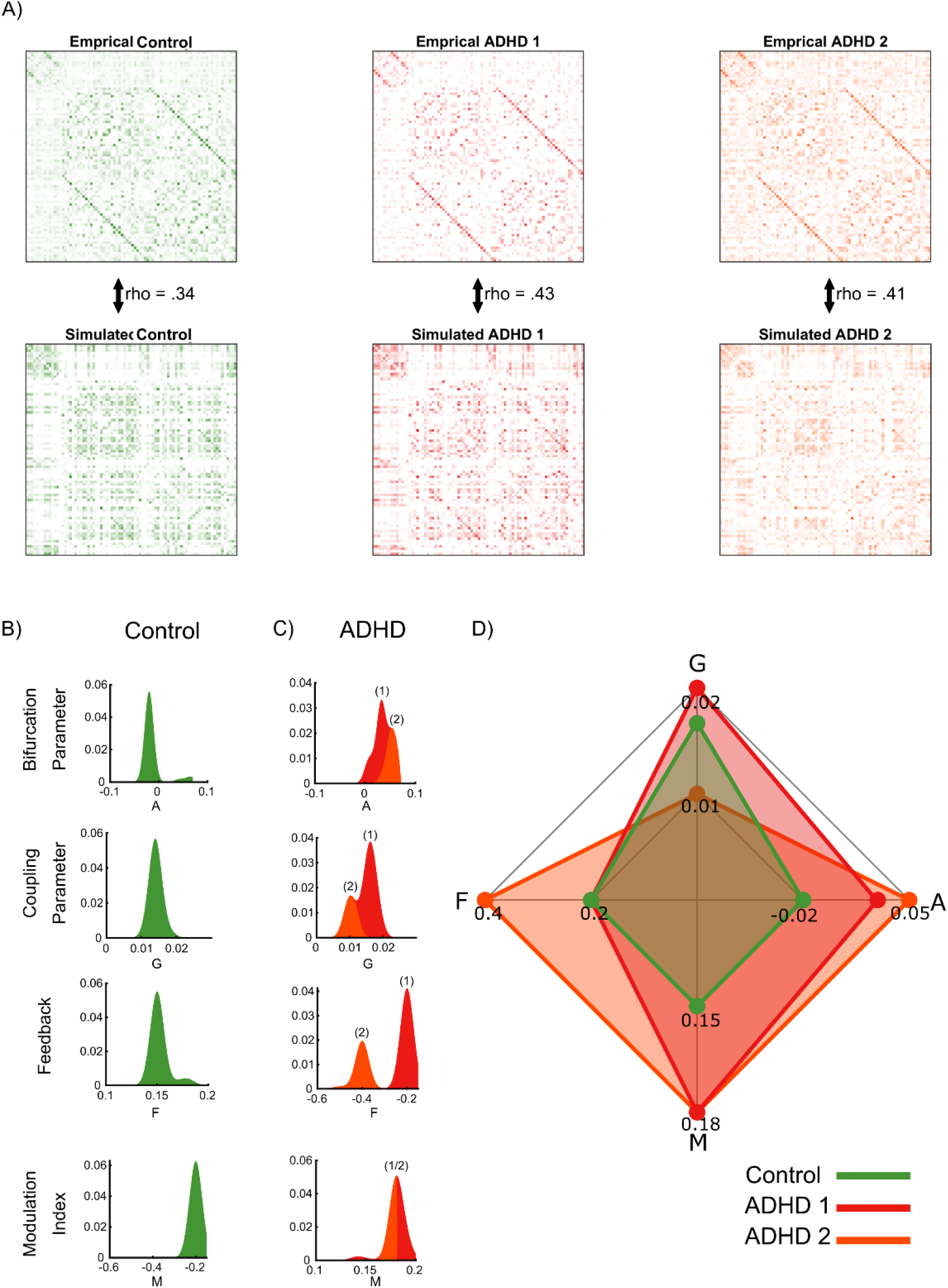
Estimated model parameter sets for ADHD and Control cohorts. (A) Empirical and simulated functional connectivity matrices for the three cohorts (Control, ADHD1, ADHD2,). The Pearson correlation values between empirical and simulated functional connectivity for the Control, ADHD1 and ADHD2 were rho = 0.34, 0.43, and 0.41, respectively. (B) The distribution of the bifurcation A, global coupling G, feedback F and modulation index M parameters yielded by 500 Monte Carlo simulations for the Control cohort (C) Same as in panel (B) for the two ADHD subgroups. (D) The parameter values for the optimal working point of the model are illustrated separately for the Control and ADHD groups.

Moreover, since we here used two different datasets obtained on different scanners, we also assessed the possible nuisance bias of scanner on our results by initially comparing the degree of similarity in the healthy cohorts taken from the two datasets. We found a high degree of correlation, rho = 0.82 (Supplementary Figure S1), between empirical FC matrices from the data obtained in the HCP and UCLA heathy cohorts. Notably, using the parameter set derived from Monte Carlo simulations based on the HCP data cohort, our model produced FC patterns that was similar to the empirical FC obtained from UCLA cohort with an even higher degree of similarity (rho = 0.40, Supplementary Figure S1). Taken together, these results suggest that bias on our results from the type of scanner used is minimal.

Specifically, using the results from 500 permutations we broke down the distribution into the four model parameters (these can be found in **Figure 2B** for Control and in **Figure 2C** for the two ADHD subgroups). The radar plot shown in **Figure 2D** highlights the differences for the four model parameters between the three groups. Specifically, the global bifurcation parameter (A) was estimated to be marginally larger for the ADHD subgroups (A_ADHD1_ = 0.0320, A_ADHD2_ = 0.0540) compared to the Control cohort (AControl = −0.02), p < 0.070). Similarly, the Modulation Index model (M) parameter was marginally larger in the ADHD subgroups than Control (M_ADHD1_ = 0.18, M_ADHD2_ = 0.18, MControl = 0.15, p < 0.09). The exact p-values were estimated as the number of the permutations that the parameter was larger in the ADHD compared to Control to the total number of permutations. Critically, we found difference between the two ADHD subgroups for the feedback parameter (F) and for the global coupling parameter (G). The feedback parameter was found to be relatively lower in the ADHD1 compared to the ADHD2 subgroup (F_ADHD1_ = 0.02, F_ADHD2_ = 0.04, F_Control_ = 0.02), whereas the global coupling parameter was found to be relatively higher in the ADHD1 group (G_ADHD1_ = 0.016, G_ADHD2_ = 0.010, G_Control_ = 0.014).

### Comparison of the empirical and simulated global BOLD signals

So far, we have shown that the sets of model parameters that maximized the similarity of FC between simulated and empirical data differed between Control and the two ADHD subgroups. Before we go on to compare amplitude differences in local dynamics, we first wanted to assess whether the simulated data and the empirical BOLD data behave similarly in the frequency domain. To recapitulate, each brain region was modeled as an oscillator with 2 state variables (Z and w) and we extracted the regional BOLD-like activation of each ROI by resampling and low-pass filtering of the real part of the first state variable (Re{Z}) (Kaboodvand et al. 2019). Moreover, the first state variable (Z) is a complex value that represents the amplitude dynamics, with the real part serving as the BOLD-like activity and imaginary part as the internal hidden state of the oscillator. We computed the power distribution for each ROI using the Welch method and then averaged across all ROIs. The results are displayed in **Figure 3A** which shows the simulated signals by our model together with the frequency amplitude spectrum shown in **Figure 3B**. As shown in **Figure 3B**, majority of the simulated signal power resides in the frequency range 0.02 to 0.12 Hz for all three groups. Similarly, the averaged power spectrum of the empirical BOLD signals across all three groups (**Figure 3C**) shows a similar presence of slow dynamics with a concentration of a signal power in the 0.02 to 0.12 Hz range. However, the full frequency bandwidth is not completely occupied leaving some room for future improvement of the model.

**Figure 3.**
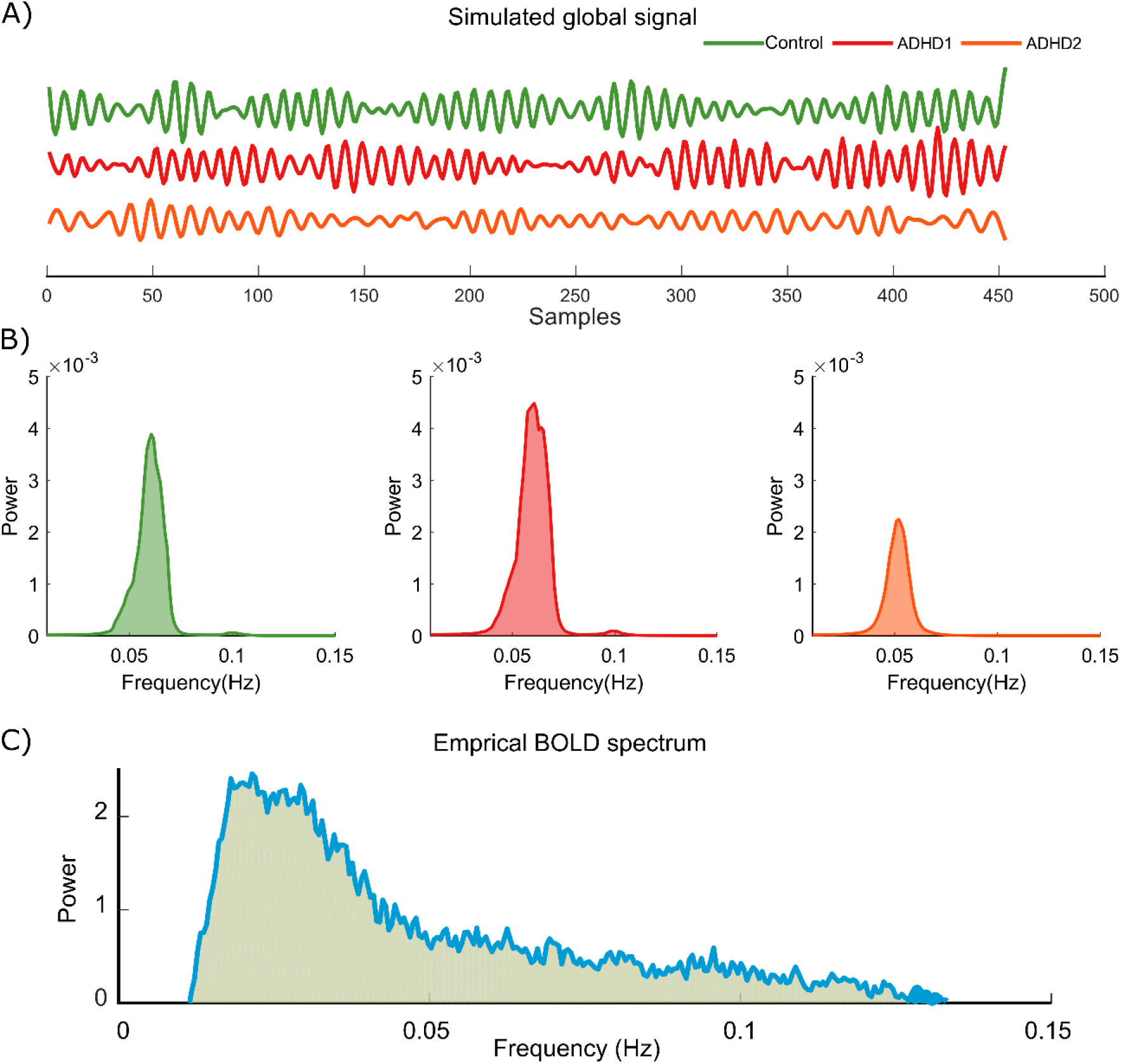
fMRI activity (global BOLD signal) and power spectrums for simulated and empirical data. (A) Simulated global signals constructed by our model using parameters sets obtained for the Control, ADHD1 and ADHD2 models. (B) The power spectrum for all three groups showed a distribution similar to the empirical fMRI data shown in panel (C).

### Determinism and entropy

As described above, the iterative tuning of parameter values converged into three parameter sets, one for the Control cohort and two for the ADHD cohort. In order to illustrate how the three parameter-sets influence the temporal dynamics of the coupled oscillators in our model, we first considered the simple case of an isolated pair of oscillators that are symmetrically and undirectedly connected to one another. The level of coupling was set to the median of the empirical SC (0.0051). Using simulated BOLD signal time-series, we numerically computed the output of the coupled oscillator model when applied to all three sets of parameters estimated in the Monte Carlo framework. The results for an oscillator of the isolated pair in the form of temporal trajectories in state-space are shown in **Figure 4**, which suggests major qualitative differences across all three groups.

**Figure 4.**
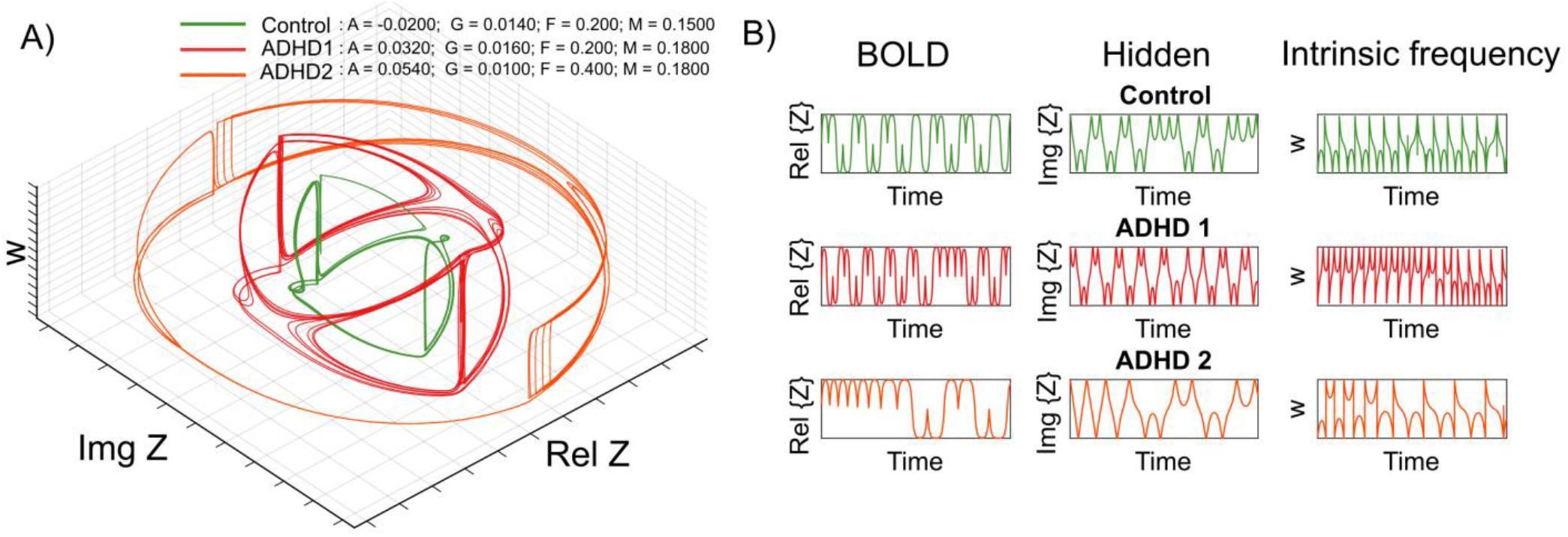
The oscillator model dynamics for the Control and ADHD subgroups. (A) An illustration of the statespace attractors of a single oscillator for the Control and ADHD models, given an isolated pair of coupled oscillators. (B) The time-courses for the simulated BOLD signal (Rel{Z}), the hidden state variable (Im{Z}) and the intrinsic frequency variable (w) are here depicted for one brain region simulated by the models for the Control and ADHD subgroups.

Nevertheless, this qualitative difference is derived from only a pair of coupled oscillators in an unrealistic arrangement in so far that they are isolated from the rest of brain network, since a full assessment of the state-space at whole-brain scale is non-trivial in this kind of analysis. Accordingly, we assessed the state space at the whole-brain scale using another approach, namely recurrence quantification analysis. Deterministic dynamical systems revisit pervious states (within an arbitrary close distance) at some later point in time (Poincaré 1890). Based on this fundamental property, assessing the recurrence patterns of a system could potentially reveal valuable information about the governing dynamics. To this end, we noted that the difference in parameter sets indicated principal qualitative differences in the dynamics between the three groups. To assess these differences further, we quantified the qualitative changes in state space using recurrence quantification analysis (Kaboodvand et al. 2020). In line with previous studies, which have shown that nonlinear signal features such as entropy can be used to characterize ADHD (Gómez et al. 2013; Sokunbi et al. 2013), we employed recurrence quantification analysis to probe the degree of Shannon entropy of the probability distribution of the diagonal line lengths and determinism across the three groups as observed from our model. Determinism which indicates the predictability of the system, was measured by computing the ratio of recurrence points that form diagonal lines to all the observed recurrence points (Marwan et al. 2007). Statistical significance was tested using 1000 iterations of Fourier-based surrogated data (Kaboodvand et al. 2020). In order to produce surrogated data, Fourier transform was applied to the original data, and for each iteration the phase content was replaced with a random phase sequence that ensured us that the generated surrogated data had a similar spectral power profile as the original data. The results in **Figure 5** show that the dynamics of the ADHD2 subgroup had a smaller degree of determinism (*CI* = [8.23, 12.09]; **Figure 5A**) and entropy (*CI* = [0.33 1.30]; **Figure 5B**) compared to both Control and ADHD1 groups. In contrast, despite showing qualitatively different state-space attractors as depicted in **Figure 4**, the dynamics of ADHD1 group showed no difference in determinism, *CI* = [-2.00, 1.14] nor entropy, *CI* = [-0.24, 0.54] compared to the Control or ADHD2 group (**Figure 5A, B**).

**Figure 5.**
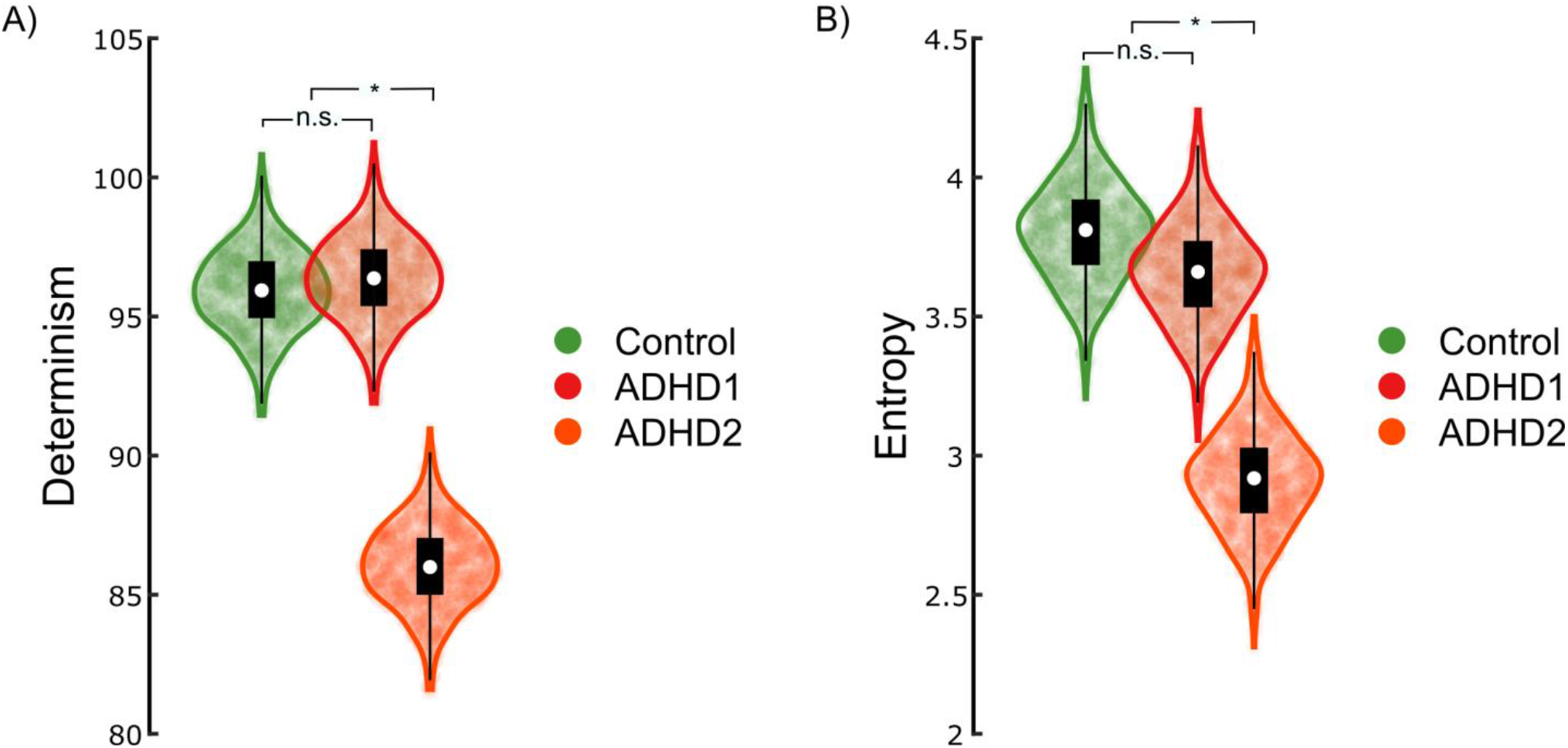
Differences in dynamical regimes across groups in terms of determinism and entropy. (A) The ADHD2 subgroup showed less determinism compared to both the Control group and the ADHD1 subgroup. (B) The ADHD2 subgroup showed less entropy compared to both Control and the ADHD1 subgroup. Each violin plot shows the distribution of 1000 surrogate data, the white dot shows the median and the scatter plot shows values of surrogated data. Interquartile ranges are shown by Whiskers.

Next, we assessed the quantitative regional changes in the amplitude of brain resting-state fMRI signals. The simulated regional BOLD amplitude, estimated by taking the mean centered root mean square (RMS), were compared across the three groups by performing pair-wised t-tests. The results are shown in **Figure 6** which shows a stronger activity in the left isthmus cingulate (*t*(81) = 3.30, *p* < 7e-4) and left lingual (*t*(81) = 3.78 *p* < 4e-4) for the ADHD1 subgroup compared to the Control group. However, the amplitude for the right pars triangularis (*t*(81) = −2.88, *p* < 2e-3) was stronger in the Control group compared to the ADHD1 subgroup (see also **Figure 6A**). For the ADHD2 subgroup versus the Control group comparison, we found similar differences in amplitude. That is, higher activity in the left isthmus cingulate (*t*(81) = 5.11, *p* < 1e-6), left lingual (*t*(81) = 3.12, *p* < 1e-3) and a trend for reduced amplitude in the right amygdala (*t*(81) = −1.54, *p* < 0.06) (see also **Figure 6B**). Moreover, for the ADHD1 compared to the ADHD2 subgroup (see also **Figure 6C**), we found significant increases in amplitude in the right amygdala (*t*(81) = 2.34, *p* < 0.01), but a decrease in the right putamen (*t*(81) = −3.34, *p* < 6e-4) and the right insula (*t*(81) = −3.18, *p* < 1e-3).

**Figure 6.**
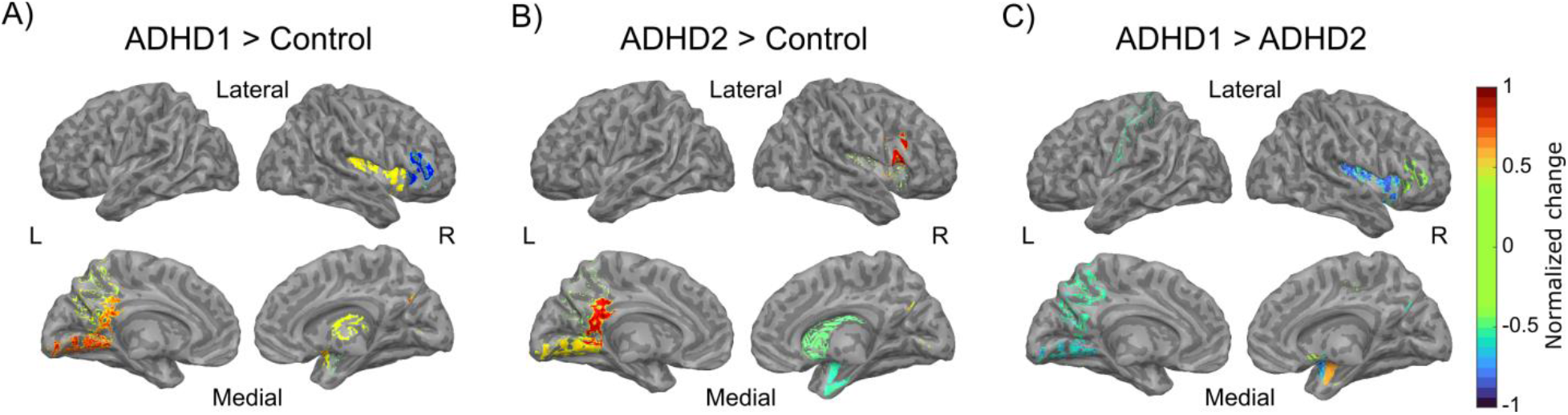
The simulated regional BOLD amplitudes differ in ADHD cohorts and Control. (A) Simulated BOLD signal amplitudes have a larger value in the left isthmus cingulate and left lingual but less value in the right pars triangularis for ADHD1 compared with Control cohort. (B) Similarly, in the simulations of ADHD2 subgroup, larger amplitudes in the left isthmus cingulate and left lingua were detected, but only marginally smaller amplitude in the right amygdala. (C) A comparison of the simulations for two ADHD subgroups revealed larger amplitude in the right amygdala but smaller amplitude in the right putamen and right insula for ADHD1 compared with the ADHD2 subgroup.

### Relationships between behavior and brain dynamics in ADHD

So far, we have presented evidence suggesting that the ADHD cohort has a heterogeneous nature from whole-brain, non-linear dynamics perspective. Based on their marked differences in BOLD signal evolution over time, our findings suggest that the functional neuroimaging data from the investigated ADHD group can be divided into two subgroups. The suggested dichotomy for the ADHD cohort based on signal dynamics begs the question of which behavioral traits that might play a significant role in this context. Obviously, there are an abundance of cognitive, sensory, emotional factors that may be relevant in this context. As noted in the introduction, recent hypotheses indicate that the heterogeneity within the ADHD cohort could be potentially manifested as differences in dysfunctional regulation of emotional information. Hence, to address this issue, we investigated whether our grouping of the ADHD cohort can also be traced to the behavioral data related to both task-related (non-emotional) and trait parameters (emotional) that were simultaneously collected for the ADHD group. A mixed effect model with groups (i.e. ADHD1 and ADHD2), age and gender as the fixed-effect terms and by-participant intercept as the random-effect term was used to predict the trait and task-related measures. We used the Stroop and the Stop-signal tasks as measures of the subject’s response inhibition ability and the results are shown in **Figures 7A** and **7B**. We found no significant difference between ADHD subgroups in terms of performance for the selective attention task estimated as the mean reaction time for incongruent trials in the Stroop task (*t*(36) = −1.69, *p* = 0.10, **Figure 7A**). Moreover, the Stop-signal reaction time (SST), which was calculated by subtracting the average Stop-signal delay from quantile reaction time, did not show a significant difference in performance between two the ADHD subgroups (*t*(36) = 1.40, *p* = 0.16, **Figure 7B**).

**Figure 7.**
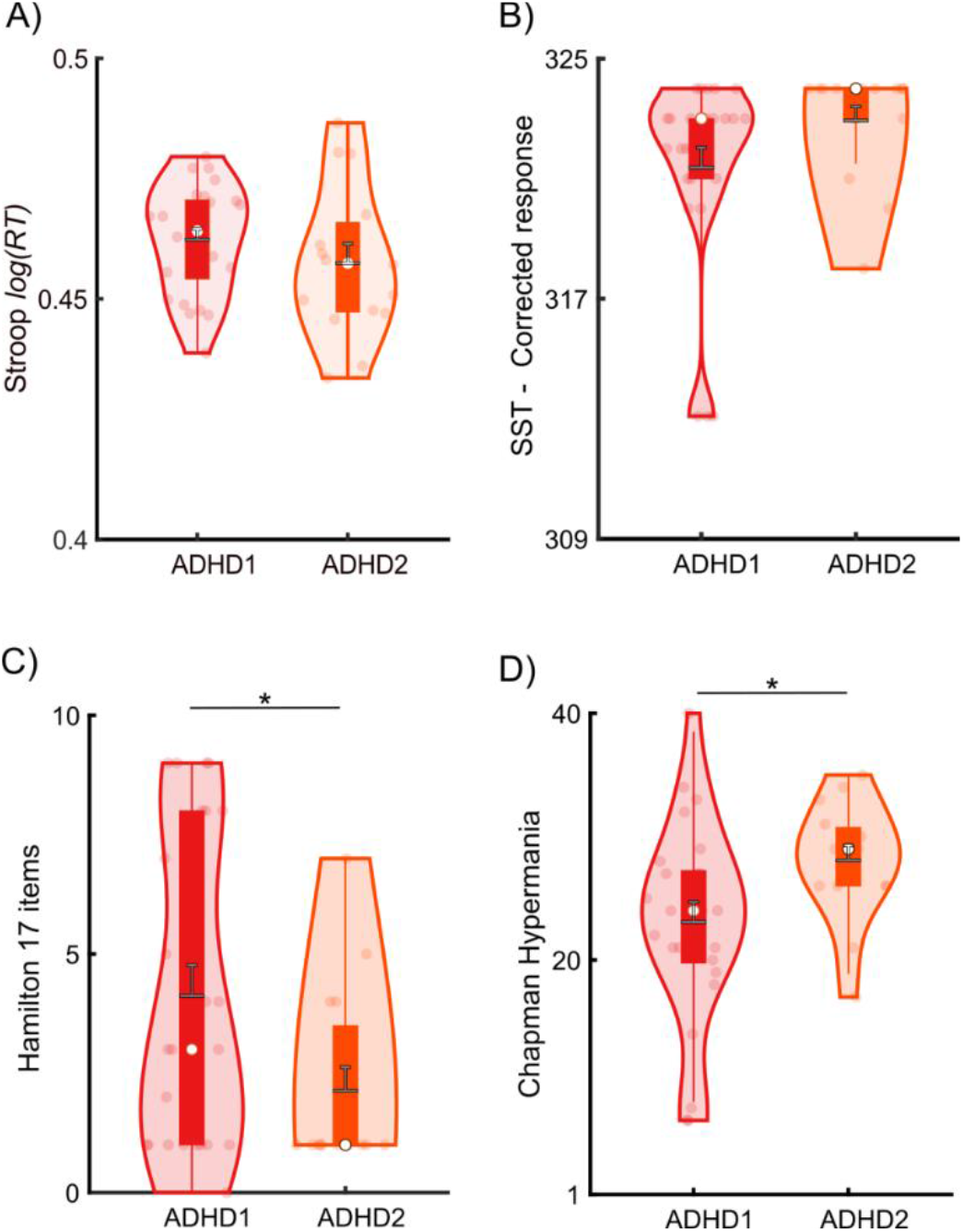
Comparison of task-related performance, mood and personality traits between the ADHD-subgroups. Violin plots are shown for two task-related measures (panels A and B) and two personality traits (panels C and D) across ADHD subgroups. The solid box indicates 25th and 75th percentiles, the white dot displays the median value, the horizontal gray line with the error bar denotes mean ± standard error. The whiskers show the maximum and minimum values within 1.5 times the interquartile range. The shaded area shows the distribution of data and each dot represents an individual data point. (A) The logarithm of reaction time for incongruent color-word trials in the Stroop task indicated no difference in performance between subgroups. (B) No significant difference between ADHD subgroups was found for the number of correct responses in the SST task. (C) A significant difference was found for the depression scores (Hamilton psychiatric rating scale) between ADHD1 and ADHD2. (D) Personality traits using the Chapman hypomanic score showed significantly differences between the two subgroups.

In contrast, trait parameters such as mood and emotion were significantly different between the two ADHD-subgroups. Specifically, the 17-item Hamilton psychiatric rating scale for depression (Hamilton 1960) and the hypomanic personality scale (Eckblad and Chapman 1986) demonstrated that the ADHD1 subgroup had higher depression scores compared to the ADHD2 subgroup (*t*(36) = −2.52, *p* = 0.016; **Figure 7C**). In contrast, the ADHD2 group scored significantly higher on the hypomanic personality scale compared to the ADHD1 subgroup (*f*(36) = 2.21, *p* = 0.033; **Figure 7D**).

### Modulating ADHD brain networks by *in silico* nodal excitatory stimulation

Finally, we assessed if in silico excitation of individual regions of the whole-brain model can change whole-brain dynamics such that the network organization of the weakly coupled oscillator model of the ADHD can move closer to the dynamics that is expressed in the Control group. For each stimulation target (82 regions), whole-brain dynamics were simulated for the ADHD subgroups and network measures were computed and normalized across simulations to construct a feature space as previously described. The results are shown in **Figure 8**. We found that *in silico* stimulation of the left medial orbitofrontal cortex (*t*(81) = 2.16, *p* < .017) and right posterior cingulate cortex (*t*(81) = 1.76, *p* < 0.04) reduced the distance between the ADHD1 subgroup and the Control cohort (**Figure 8A**). Moreover, for ADHD2 model, we observed that excitatory stimulation of left rostral middle frontal (*t*(81) = 2.22, *p* < 0.015) and right frontal pole (*t*(81) = 2.40, *p* < 0.01) decreased the distance to the Control cohort (**Figure 8B**).

**Figure 8:**
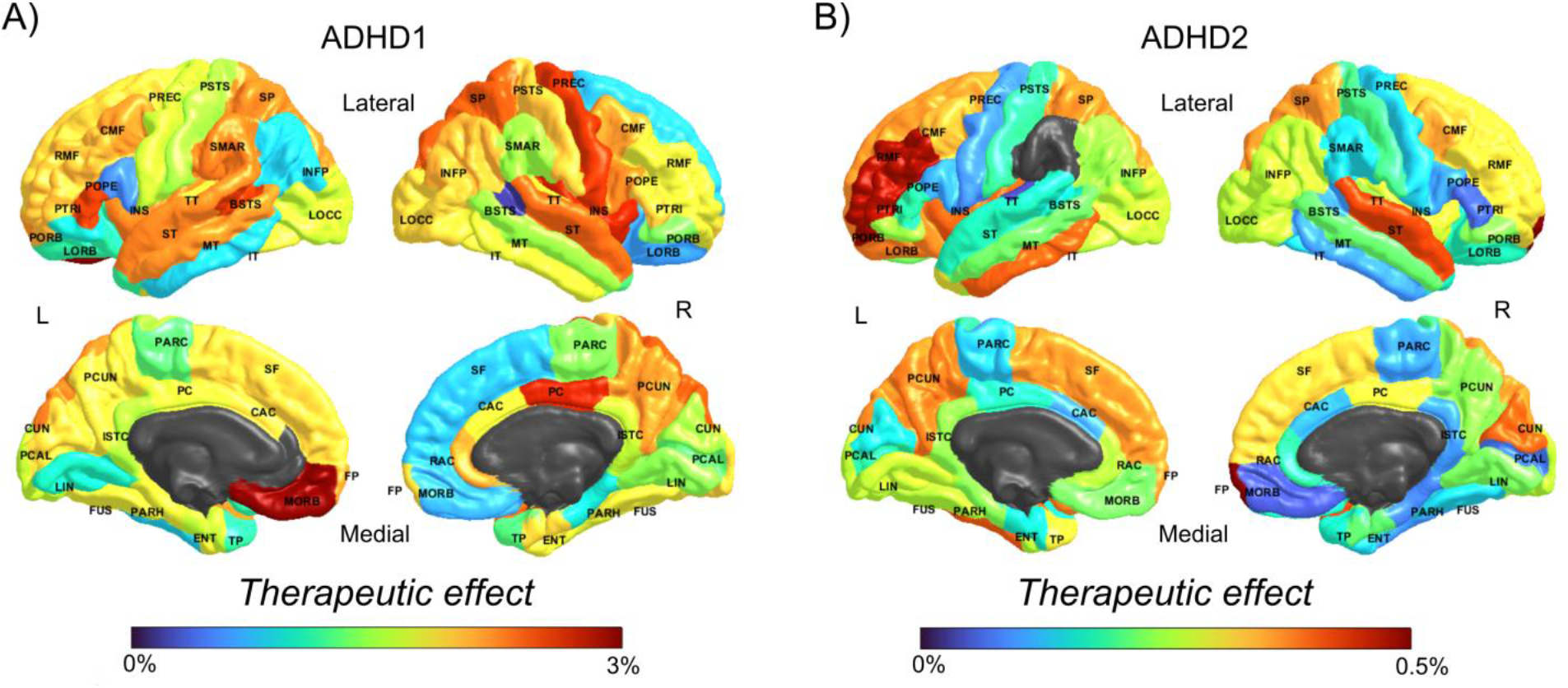
The effect of nodal in silico excitatory stimulation. (A) The excitatory therapeutic map of ADHD1 showed left medial orbitofrontal and right posterior cingulate as the most effective stimulation sites. The effect sizes for all cortical regions are shown in color coded. (B) The excitatory therapeutic map for ADHD2 showed the left rostral middle frontal and right frontal pole as the most effective stimulation targets.

We assessed the effects of the intensity of simulated brain stimulation for only ADHD1 cohort, where we found non-negligible effect on the group level. We found a negative relationship between the intensity of stimulation (parameter A in our model) and the Euclidean distance in the medial orbitofrontal region (slope = −2.97, Supplementary Figure S2). The Euclidean distance was calculated on the relative difference of the centrality measures estimated in the stimulated ADHD and the Control cohort models. As a control analysis, we also assessed the banks of superior temporal sulcus, a region that our model predicted to not be of therapeutic value in ADHD1 cohort by means of external excitation. For this specific region, we found a positive reverse relationship (slope = 6.19) with stimulation intensity (Supplementary Figure S2).

## DISCUSSION

We used a whole-brain, non-linear dynamics modeling approach to assess differences between a Control and ADHD cohorts in resting-state fMRI brain dynamics. The results from our simulations suggest that the investigated ADHD cohort can, based on quantitative differences in the dynamics, be divided in to two subgroups, each with their own personality traits. Importantly, we demonstrated that the two suggested subgroups of ADHD captured both functional as well as behavioral differences. Several morphological studies have shown that there is a global reduction in gray matter volume in the ADHD brain (Carmona et al. 2005; Nakao et al. 2011). In addition to the structural changes, pervious fMRI studies have confirmed an overall elevation in brain activity in ADHD (Tian et al. 2008), possibly due to compensatory mechanisms to mitigate reduction in gray matter density. In the present study, we found that the global bifurcation parameter was higher in both ADHD subgroups compared with the Control group, suggesting that the local dynamics of the oscillators in the ADHD brain evolve over time with elevated amplitude compared with that of the healthy brain. On the other hand, it has also been hypothesized that impairment in the mechanisms that govern neural gain may play a role for ADHD-like neurocognitive impairments (Hauser et al. 2016). Interestingly, we also found that the global coupling parameter was lower in ADHD, but only in one of the subgroups, namely ADHD2. This finding suggests that the neural gain impairment often exists as a deflection in the ADHD brain, but it may not always be present. The lower gain in the ADHD2 subgroup, together with elevated bifurcation parameter caused a major shift in the dynamics of the simulated ADHD brains by displaying less determinism and entropy. This observation can possibly be linked to the increased unpredictability across various cognitive domains typically found in ADHD (Hauser et al. 2016). Notably, a previous study found less entropy or complexity in ADHD patients and showed negative correlation with symptom scores (Sokunbi et al. 2013).

Of note, there was a higher activity predicted for the left isthmus cingulate cortex (also known as retrosplenial cortex) for both ADHD subgroups compared with the Control group. It was previously shown that the retrosplenial cortex serves as the provincial hub of the default mode network (DMN) and it mediates functional interactions (integrates information transfer) among DMN subsystems (Kaboodvand et al. 2018). Moreover, our previous study found the most commonly recruited configuration of FC patterns, associated with segregated DMN sub-systems, was employed for a shorter duration by the ADHD cohort compared with the Control group (Kaboodvand et al., 2020). Our finding of ADHD-related hyperactive retrosplenial cortex in this study well explains our previous observation regarding the over-integration of the DMN sub-systems for the ADHD subjects. There is also evidence regarding the critical role of retrosplenial cortex in the whole-brain dynamics and high level of hazardousness for this region, in a sense that perturbing the retrosplenial cortex inflicts considerable damage on the whole-brain network dynamics (Kaboodvand et al., 2019).

Previous studies have reported altered structural morphology and resting state FC of the insular cortex in ADHD (Zhao et al. 2017; Lopez-Larson et al. 2012). Presumably, insular cortex is a key region of the salience network (Uddin 2015) and it has been linked to impairment of executive functions in ADHD (Zhao et al. 2017). In addition to cortical regions, meta-analysis suggests involvement of subcortical regions as well (Cortese et al. 2016). Putamen is thought to be involved in the cortico-striatal circuit that has been linked to ADHD pathology (Bush et al. 2005). Moreover, decreases of gray matter has been shown in right putamen in ADHD (Ellison-Wright et al. 2008). Interestingly, our findings suggest that the ADHD subgroups have different levels of activity in the right amygdala, right insula and right putamen. Importantly, we showed that the ADHD subcategorization according to our model was mirrored in the empirical behavioral data with significantly different depression and hypomanic personality trait scores; a fact possibly explained by the difference in amygdala, putamen, and insula cortex activities (Cullen et al. 2014; Guo et al. 2012; Sprengelmeyer et al. 2011). These findings directly support the use of a dimensional approach to categorize ADHD, with ADHD phenotypes emerging and differentiating according to dimensions of emotional instability (Petrovic and Castellanos 2016). Our findings support the notion that at least two pathological dynamics are involved in ADHD that could be tied to emotional traits.

We investigated if our model could serve as a guide for a therapeutic use of excitatory stimulation across the ADHD spectrum. Applying focal brain stimulation (e.g. repetitive TMS) has been shown to selectively change the oscillatory activity of the targeted region and modulate the frequency and power of oscillations (Thut and Pascual-Leone 2010; Okamura et al. 2001). There are previous efforts for modelling the frequency effects of neurostimulation (Gollo et al. 2017; Cocchi et al. 2016), suggesting its applicability in rectifying atypical intrinsic timescales as observed for example in autism disorder (Watanabe et al. 2019; Gollo 2019). Relatedly, in this study we observed ADHD-related differences in time-varying intrinsic frequency dynamics which is possibly a notion of impaired intrinsic time scales in ADHD. Nevertheless, in the current study we only focused on the amplitude dynamics and resting state FC.

We found that the left medial orbitofrontal and the right posterior cingulate cortices were the most beneficiary stimulation sites in ADHD1, whereas the left rostral middle frontal and right frontal pole were the best candidate targets for the ADHD2 subgroup. A possible biological route to achieve a therapeutic use of the predicted effect on brain dynamics from external excitatory stimulation is the modulatory effect provided by TMS that is believed to affect hypo-functional dopamine terminals and thereby yield a local “boosting” in neuronal dopamine signaling. However, the effect-sizes were small in the ADHD1 subgroup (3%) and negligible in ADHD2 (.5%). The limited effectiveness of externally given stimulation as reported here emphasizes the notion that the localization of stimulation targets cannot be based solely on separating ADHD patients into two sub-groups. Rather, it is likely that larger effects sizes will require individualized stimulation protocols. Moreover, in future studies it would be worthwhile to investigate whether the effect sizes from external stimulation can be increased by using a finer spatial granulation in the analysis, e.g. the Schafer-Yeo parcellation that allows for up to 1000 parcels (Schaefer et al. 2018), or by moving from region-based analysis to cortical surface-based modelling while incorporating local connectivity into the wholebrain model (Coombes et al. 2012; Spiegler and Jirsa 2013). It is therefore perhaps not overly surprising that previous attempts to apply TMS for the purpose of alleviating symptoms in ADHD have failed to show any significant therapeutic effect (Weaver et al. 2012). However, individualized whole-brain modelling can serve as a useful tool for exploring the large parameter space (e.g., various stimulation targets) in order to achieve a well-optimized stimulation protocol and improve TMS efficacy.

In general, dynamical system modeling approach used here allows for investigations of neural dynamics that are less biased by inter-individual variation and experimental confounds such as headmotion which has been shown to have detrimental effect on resting-state fMRI FC analyses (Power et al. 2018). This has been achieved by combining the multimodal imaging and confining the whole-brain dynamic behavior by the local dynamics (Kaboodvand 2019). Therefore, the detrimental effect on resting-state fMRI connectivity analysis from motion is to some degree mitigated by the computational modeling approach that converges both structural and functional connectivity in a single mathematical framework (Kaboodvand 2019). However, one should also be aware that the correspondence between structural and functional connectivity is not straightforward. It has been estimated that the correlation between structural and functional connectivity is only between 0.3 and 0.7 (Suárez et al. 2020). Although, our model’s correlation between simulated and empirical data reached to the value that is above the typical intrinsic correlation between the SC and FC, this value is modest, and the findings should be interpreted with caution.

We did not use cohort-specific SC, but rather a universal SC obtained from HCP 500-subject release in our whole-brain simulations to minimize the potential errors due to construction of SC in an undersample dataset. Notably, the high quality and large sample size of the structural HCP dataset limits the influence from the potential spurious connections in the individualized tractography analysis. Although the anatomical structure of the brain has a critical role in whole-brain modelling, the more accurate methods of delineating SC construction are not applicable in humans. On the other hand, applying tractography to the diffusion weighted imaging (diffusion tensor imaging (DTI) and diffusion spectrum imaging (DSI)) for non-invasive reconstruction of the white-matter tracts and constructing the SC, despite being the most common practice, is not without controversy in under sampled datasets or at the individual data level. For example, comparing the diffusion MRI-based tractography with neuroanatomical tracer studies in macaque indicated above chance performance but far from perfect (Donahue et al. 2016). Therefore, the superiority of using individualized SC for whole-brain modelling is questionable. Last but not least, previous studies have not found any significant ADHD effect on the tractography-based SC (Batty et al. 2010; Carmona et al. 2005). Overall, the ADHD-related difference in the SC, particularly with the applied low resolution (i.e., 82 regions) seems to be insignificant and it further supports the idea of using a universal SC for both Control and ADHD groups. This also guarantees that the observed differences in the brain dynamics are not simply driven by the spurious differences of underlying connectomes.

We restricted our neuro-stimulation modelling to induce the oscillatory behavior in different targeted regions, to further elaborate the heterogeneity within an ADHD cohort and highlight the importance of designing stratified treatment. That being said, in future work, a spectrum of local neural excitatory and inhibitory effects (e.g. induced by different TMS protocols) combined with a more detailed and refined model, including modelling of biophysical (Efield modelling) and neurophysiological (e.g. neuroplasticity) effects, might be considered. Thus, the present work constitutes only an introductory study to highlight the importance of stratified treatment and the potential benefits of whole-brain dynamic modelling.

Finally, stimulation targets found to be of interest by the analysis described in this study, especially frontal and subcortical regions, may not be feasible to stimulate with techniques such as TMS. Nevertheless, there are other alternative stimulation technologies available that are able to target deep and frontal structures, for example using temporal interfering electrical fields (Grossman et al. 2017). Yet, we leave a more comprehensive and detailed comparison between different neuro-stimulation effects (including both changes in the local neuro-dynamics and effects propagated across the whole-brain effective connectivity network) for future work. To the best of our knowledge this has not been done yet.

In summary, we have shown that our computational modelling approach could to some extent replicate spontaneous fMRI activity in both cohorts. Specifically, our simulations and comparisons with empirical data suggest that there are good reasons to believe that the heterogeneity of symptoms and behavioral traits that are typically observed in ADHD patients can be traced back to the dynamics of resting-state fMRI signals. The observed heterogeneity of the ADHD cohort together with the different in silico excitatory stimulation effect maps, highlight the importance of designing stratified neuro-stimulation protocols. The results presented in the current study is based on relatively small cohorts and only a limited sample of the full spectrum of behavioral parameters that are typically recorded in studies of ADHD were included in the analysis. Moreover, the similarity between the empirical and simulated FCs is modest and the findings should be interpreted with caution. Therefore, future studies of brain dynamics in ADHD are needed in order to assess the generalizability of the current results in larger cohorts that includes a larger span of behavioral scores and parameters. Finally, we have presented promising results that pertain to the possibility to investigate the feasibility of in silico excitatory simulations that may serve as a theoretical basis for focal brain stimulation protocols (e.g. TMS) in patients diagnosed with ADHD.

## Acknowledgments

Data were provided by the Human Connectome Project, WU-Minn Consortium (Principal Investigators: David Van Essen and Kamil Ugurbil; 1U54MH091657) funded by the 16 NIH Institutes and Centers that support the NIH Blueprint for Neuroscience Research; and by the McDonnell Center for Systems Neuroscience at Washington University. The funders had no role in study design, data collection and analysis, decision to publish, or preparation of the manuscript.

## Data availability

Data used in this study is publicly available at https://db.humanconnectome.org and from the University of California LA Consortium for Neuropsychiatric Phenomics study.

## Code availability

The code for analyzing the data used in this study is available from the author upon reasonable request.

