## Supplementary Figure for "Whole-brain modelling of resting state fMRI differentiates ADHD subtypes and facilitates stratified neuro-stimulation therapy"

Supplemnatry Figures

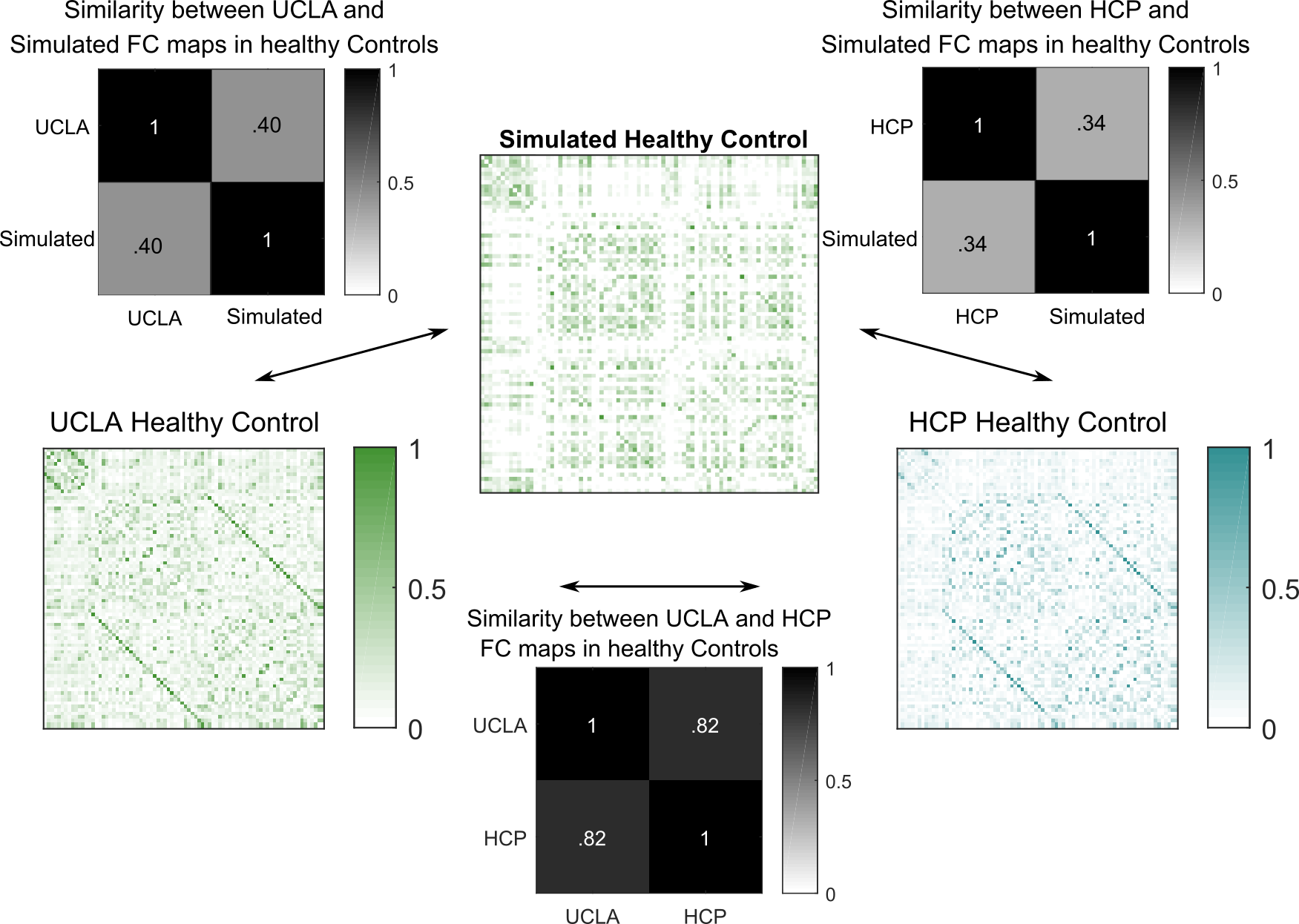

**Figure S1.** The empirical FC for the UCLA and HCP datasets showed a high degree of correlation (rho = 0.82). The correlation between empirical FC obtained from the UCLA dataset and the simulated FC matrix, given the parameter set derived from fitting the Healthy model to the HCP dataset, is even larger (rho = 0.40) than the correlation between empirical FC obtained from the HCP dataset and the simulated FC (rho = 0.34).

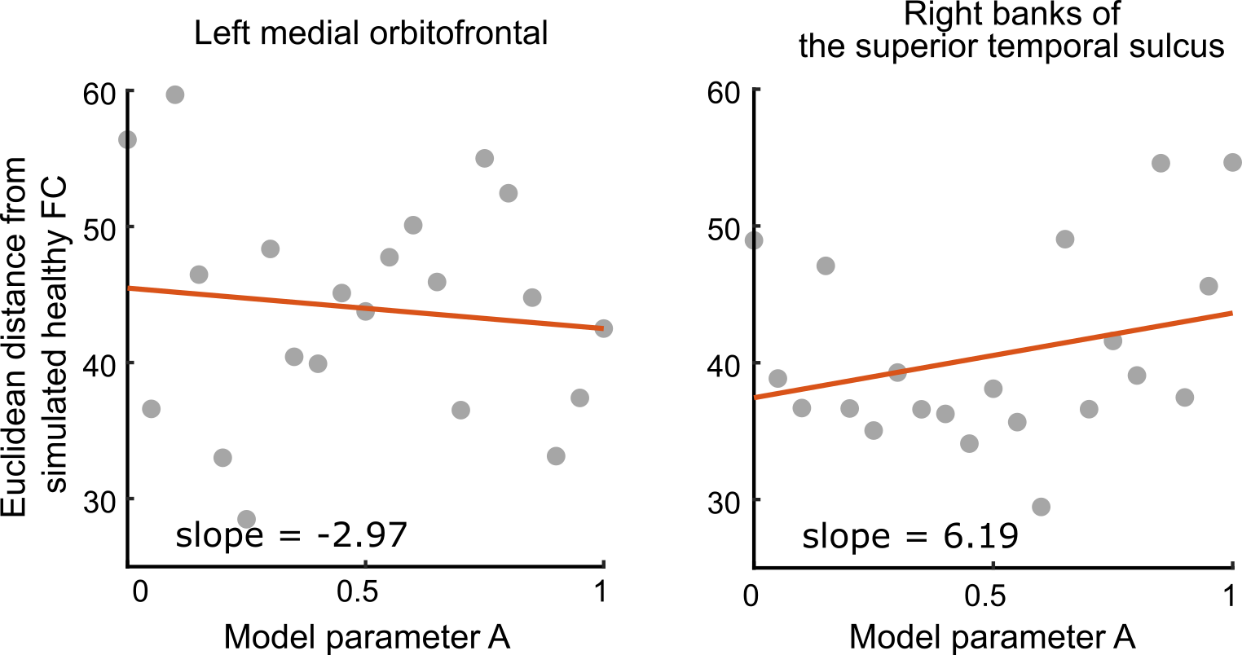

**Figure S2.** Excitatory stimulation for the left medial orbitofrontal region (the region which was found to have the highest therapeutic effect) and the right banks of superior temporal sulcus (as a control region) was repeated 20 times (with local bifurcation parameter A changing from -0.02 to 1, with the step of 0.05), to assess the relationship of the brain stimulation intensity and the therapeutic effect size. Accordingly, the Euclidean distance between the simulated ADHD1 and Heathy FC model were computed for centrality measures as a function of parameter A (representing the intensity of stimulation),where we found a negative association for the left medial orbitofrontal region and a positive association for the right banks of the superior temporal sulcus as control region.
